# ECM (collagen) mediated signaling drives the emergence of androgen independence in prostate cancer cells

**DOI:** 10.1101/2025.07.07.663456

**Authors:** Apoorva Abikar, Prathibha Ranganathan

## Abstract

The extracellular matrix is a dynamic entity that undergoes continuous deposition, remodeling, and degradation to maintain tissue homeostasis. As a major structural component of the tumor microenvironment, the ECM plays crucial roles in providing mechanical support, modulating the microenvironment, and serving as a reservoir for signaling molecules. Tissue stiffness is primarily determined by the abundance and cross-linking of the ECM components. Collagen is one of the major components of ECM and contributes to 30% of the total ECM. Increased collagen deposition contributes to enhanced tissue stiffness and is associated with a more aggressive cancer phenotype. A restructured ECM, particularly collagen content, can influence cellular signaling pathways through interaction with cell surface receptors. In the current study, we investigated the role of collagen in prostate cancer. Our findings indicate that collagen, a major component of the TME, modulates prostate cancer cell migration, alters sensitivity to chemotherapeutic agents, and promotes non-genomic activation of androgen receptor signaling through FAK activation, thereby contributing to therapy (androgen deprivation therapy) resistance and cancer progression.

## Introduction

Tumor microenvironment (TME) is a heterogenous mixture of both cellular (tumor cells, fibroblasts, endothelial cells, adipocytes, immune cells, and neuroendocrine cells) and acellular components (extracellular matrix proteins, antibodies, cytokines, extracellular vesicles, growth factors, and metabolites). The tumor behavior is not solely defined by the factors produced by the tumor itself, but also heavily influenced by the complex interaction between the tumor and its microenvironment. This interaction is what shapes the tumor microenvironment and influences cancer cell survival, immune surveillance, angiogenesis, local invasion, metastasis, and therapeutic response [1]. In the early days, tumor intrinsic factors were considered the primary reason for therapy resistance and cancer progression. However, recent research studies have demonstrated that components of the ECM and other factors within TME contribute significantly to tumor behavior and therapeutic failure (reviewed in [2–4]).

Extracellular matrix (ECM) is a 3D network of macromolecules that provides structural and biochemical support to cells [5]. Animal ECM can be classified into two main components: the interstitial matrix, which surrounds the cells, providing structural support and facilitating cell-cell communication; and the basement membrane, a thin sheet that separates the stroma from cells of various origins, playing a key role in tissue integrity and function [6]. In cancer, ECM is biochemically different and significantly stiffer than normal ECM. It undergoes extensive remodeling and contributes significantly to cancer cell proliferation, survival, metastasis, and response to therapy [5].

Collagen is one of the most abundant components of the ECM and contributes about 30% of the total body protein [7, 8]. Collagen is a structural protein that provides structural support to the extracellular space of connective tissue. Due to its rigidity and structural resistance, it serves as an ideal scaffold for skin, tendons, bones, and ligaments. Collagens are classified into various types based on the type of structures they form. The most common types are type I, type IV, with type I comprising over 90% of the collagen in the human body [9]. Collagens are typically homo or heterotrimers composed of one, two, or three different polypeptide chains [7]. Collagen is critical in shaping tumor behavior; tumor cells reverse the process by reshaping collagen to form a reinforcing cell-collagen loop. This dynamic and reciprocal interaction would gradually foster cancer progression [10]. Adhesion of COLI and COLIV to cancer cells impacts cancer progression [11].

Collagen fibres constitute the major structural component of the ECM in the normal prostate stroma. Among these, collagen type I and type III are the predominant fibrillar collagens, whereas collagen type IV is primarily localised in the basement membrane (reviewed in [12, 13]). Excessive synthesis of type 1 collagen has been observed to activate periacinar fibroblasts adjacent to prostatic intraepithelial neoplasia, a precursor lesion of prostatic adenocarcinoma. This suggests that increased collagen production within the prostate is associated with the development and progression of prostate cancer (PCa) [14, 15]. An integrative study by Xiao, Y., Lai, C., Hu, J. *et al.* reported that collagen-associated genes like PLOD3, COLA1, MMP11, and FMOD could potentially act as a prognostic marker for biochemical recurrence-free survival of patients [16]. Differential expression of collagen has been observed between benign prostate hyperplasia (BPH) and prostate cancer. For example, PLOD3, a collagen-associated gene, is differentially expressed in cancer and para-cancer prostate tissues of clinical specimens [16]. Collagen XXII protein levels are significantly elevated in prostate cancer compared to benign prostate tissue. Notably, its collagen XXII expression is higher in distant metastases or prostate cancer compared to either localized or regional (lymph node) metastases [17]. Furthermore, patients exhibiting higher levels of collagen XXII expression demonstrate a 2.8-fold increased risk of PSA recurrence. This suggests that distinct patterns of collagen expression may serve as a potential molecular biomarker for prostate cancer progression, metastasis, and patient survival [17, 18].

In our laboratory, we have previously conducted transcriptomic analysis comparing normal vs cancerous prostate tissue [19] as well as normal vs cancer-associated fibroblasts from the prostate [1]. In addition, we performed quantitative mass spectrometry analysis on the conditioned media of cultured fibroblasts (normal vs cancerous fibroblasts) and identified collagen as one of the differentially expressed factors [20]. Several collagen isoforms were found to be differentially expressed across these datasets. Building on these findings, the current study investigates the effect of collagen on prostate cancer cell migration, drug sensitivity, stemness, cell proliferation, and androgen signaling.

## Materials and methods

### Cell culture

Prostate cancer cell lines LNCaP, DU145, and PC3 were procured from the National Centre for Cell Science (NCCS), Pune, India. LNCaP cells were grown and maintained in Rosewell Park Memorial Institute-1640 (RPMI-1640) (Thermo Fischer Scientific, Cat no: 23400-021) media containing 15% fetal bovine serum (heat inactivated) (Thermo Fischer Scientific, Cat no: 10270), 1X PenStrep (Thermo Fischer Scientific, Cat no: 15140122), and 1X Glutamax (Thermo Fischer Scientific, Cat no: 35050061), whereas PC3 and DU145 cells were maintained in DMEM-F12 (Thermo Fischer Scientific, Cat no: 12400024) media supplemented with 10% FBS, 1X PenStrep, and 1X Glutamax. All cells were maintained in a humidified 37°C incubator maintaining 5% CO_2_.

### Collagen coating and treatment with growth factors and inhibitors

Lyophilized collagen (collagen 1) (Merck Roche, Cat no: 11179179001) was reconstituted using Dulbecco’s phosphate-buffered saline (DPBS) (Thermo Fischer Scientific, Cat no: 21600-010) containing 0.1% acetic acid. The reconstituted collagen was then coated on the plates or wells (5ug/cm^2^).

Following collagen coating, cells were seeded in plates/wells containing FBS and antibiotics in the media. After 6-8 hours of seeding, the cells were treated with the required amount of inhibitors or growth factors for 48hrs in serum-free media. The drug was replenished after 48hrs. The control/vehicle cells were treated with DMSO. At 72hrs, the RNA was isolated and converted to cDNA. The concentration of activators and inhibitors used is summarized in Table 1

**Table 1:** List of growth factors and inhibitors used in the current study. The list shows the drug name, catalog number, manufacturing company, and the abbreviations and concentration used in the current study

| **Drugs** | **Catalog No** | **Company** | **Abbreviation** | **Concentration** |
| --- | --- | --- | --- | --- |
| PF 573228 (FAK inhibitor) | 14924 | Cayman Chemicals | FAK inhi | 10nM |
| Flutamide | 20359 | Cayman Chemicals | Flut | 5μM |
| Erlotinib | 10483 | Cayman Chemicals | Erlo | 5μM |
| EGF | P01133 | Thermo fischer scientific | EGF | 10nM |
| Docetaxel | 11637 | Cayman Chemicals | Doce | 0.1-10nM |
| Collagen from Rat tail | 11179179001 | Merk Roche | Col | 5μg/cm2 |

### RNA isolation

Total RNA was isolated using RDP trio Reagent (Himedia, Cat No: 15596026) according to the manufacturer’s instructions. RNA was quantified on a nanodrop (Thermo Scientific™ NanoDrop™). The quality of RNA samples was assessed by running them on a 1% agarose gel.

### qRT-PCR

2µg RNA was converted to cDNA using a Verso cDNA synthesis kit (Thermo Fisher Scientific Cat no: AB1453A) according to the manufacturer’s instructions. 20ng RNA equivalent cDNA was used for the PCR reactions. qRT-PCR was carried out using the Kapa SYBR FAST Universal 2XqPCR Master Mix (Roche, Cat no: KK4601). RPL35 served as an internal control. The expression levels of each gene test sample relative to the control were analyzed using the ddCt method. Relative fold changes with respect to control/vehicle samples were plotted in the graphs.

### Trypan blue exclusion assay (Chemosensitivity assay)

0.05X10^6^ cells/ well were plated in the 24-well plate (either collagen-coated or plain surface) and allowed to attach. After 6-8 hours of incubation, docetaxel was added to each in increasing concentration (6 different concentrations of drug, including the vehicle cells). The drug was replenished after 48hrs. The control/vehicle cells were treated with DMSO. After 72hrs, the cells were trypsinized and counted using the trypan blue exclusion assay. The percentage of viable cells was calculated. For each condition, the cells were plated in duplicate wells, and each well was counted twice. The experiment was repeated 3 times, and the average has been plotted.

### Migration assay

A total of 4×10^6^ cells were plated onto a 60mm dish (either collagen-coated or plain surface) and allowed to grow until they reached a confluent monolayer. A scratch/wound was made in each dish using a 200µL micropipette tip. The width of the scratch was measured every hour using the Nikon NIS Element software in conjunction with a light microscope at 10X magnification. At each time point, three readings were taken and the average distance was calculated. The rate of migration was calculated using the formula

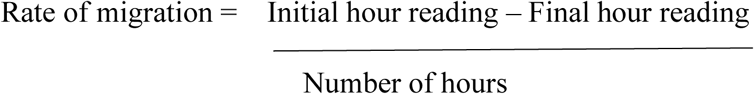

### MTT assay

5000 cells/well were plated in the 96-well plate (either collagen-coated or plain surface) and allowed to attach. After 6-8 hours of incubation, inhibitors/activators were added to serum-free media. The drug was replenished after 48hrs. At 72 hours, MTT reagent was added to the wells. After the formation of violet formazan crystals, the crystals were dissolved using laboratory-grade 100% DMSO. The experiment was repeated 3 times, and the average has been plotted.

## Results

### Expression levels of various collagen types across transcriptomic and proteomic datasets

Several collagen types were found to be overexpressed across multiple datasets, such as BPH vs PCa, fibroblasts derived from BPH and PCa, as well as the proteome derived from the fibroblasts [1, 19, 20]. These results are summarized in Table 2.

**Table 2:**
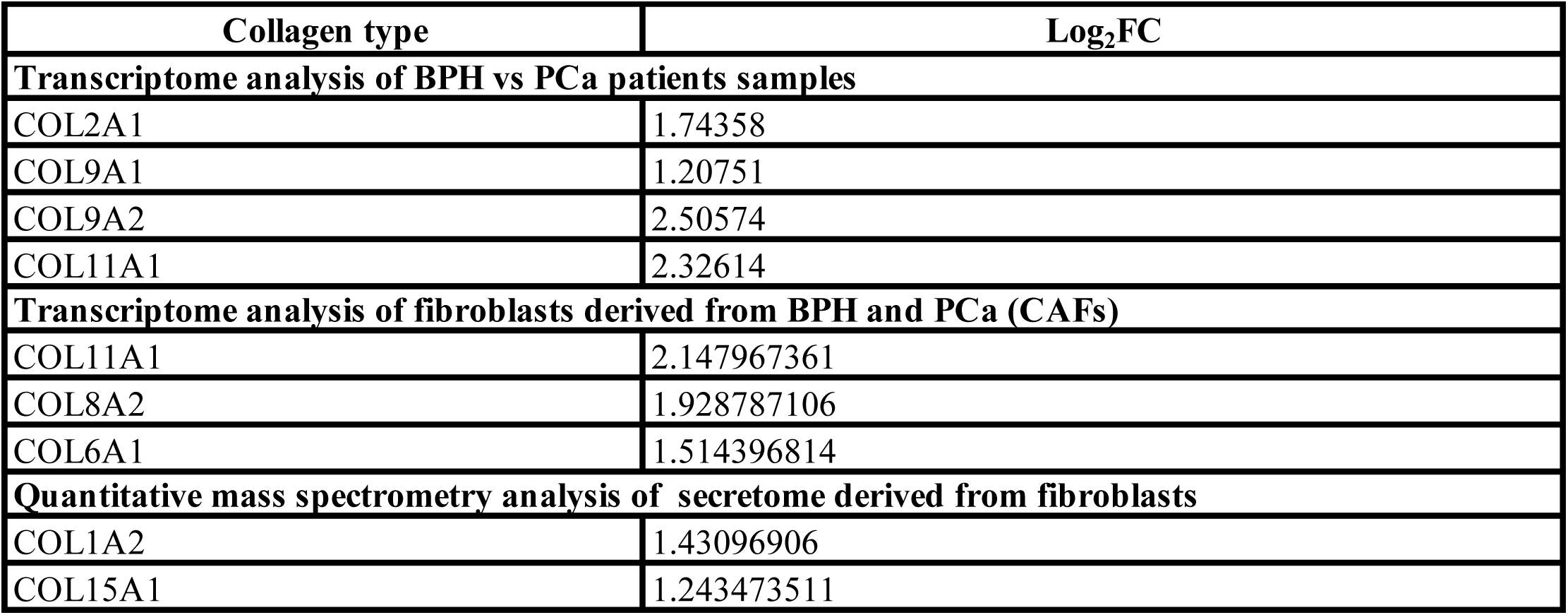
Collagen expression across different transcriptomic and proteomic datasets. The list shows the collagen type and its corresponding log_2_FC value in different transcriptomic and proteomic datasets

### Validation of collagen overexpression in CAFs

Various transcriptomic and proteomic datasets generated in our laboratory revealed the overexpression of different subsets of the collagen family. To validate these omics-based observations, we have performed qRT-PCR to assess the expression of COL11A1 RNA isolated from normal fibroblasts and cancer-associated fibroblasts. Analysis demonstrated the significant overexpression of the COL11A1 gene in CAFs compared to normal fibroblasts (Figure 1).

**Figure 1:**
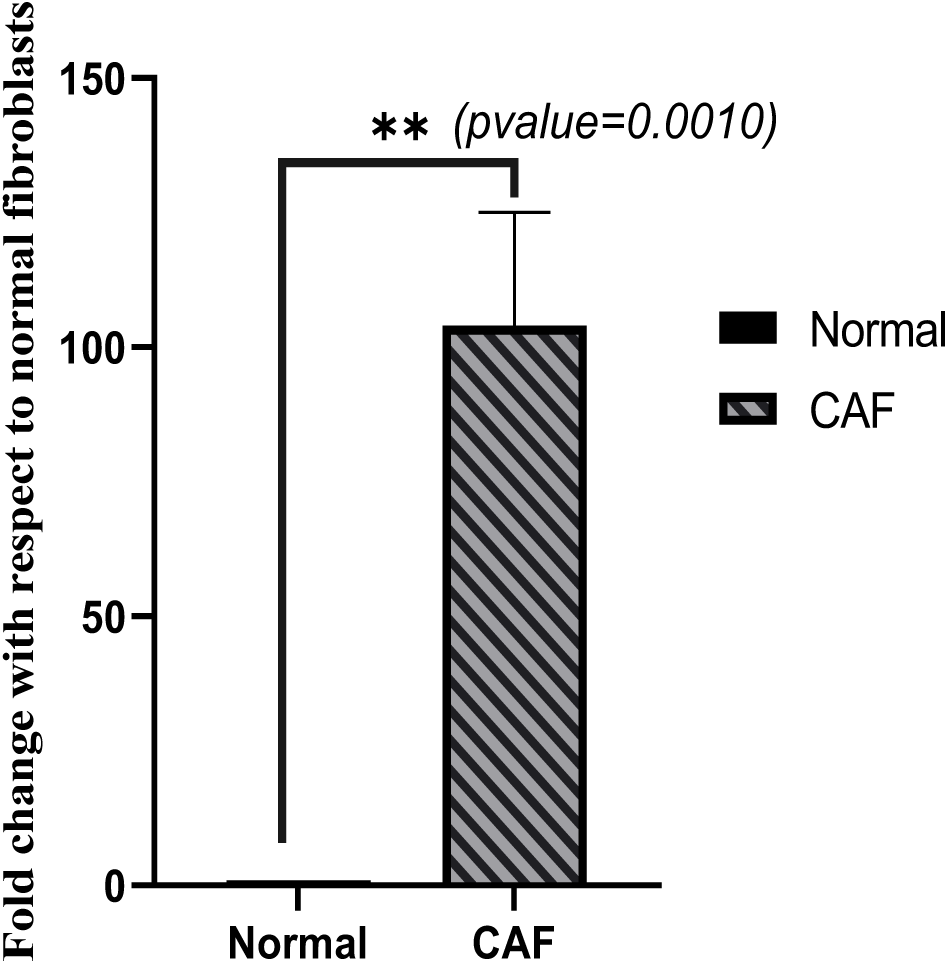
Expression of COL11A1 in cultured fibroblasts. RNA was isolated from the cultured fibroblasts and subsequently converted into cDNA, which was used as a template to perform qRT-PCR. The bar represents the relative expression levels of COL11A1 with respect to normal fibroblasts ± S.E.M., normalized to RPL35.

### Effect of collagen on the sensitivity of prostate cancer cells to docetaxel

In the current study, we evaluated the sensitivity of prostate cancer cell lines (LNCaP, DU145, and PC3) to the common chemotherapeutic drug docetaxel in the presence of collagen. The cell viability was estimated using the trypan blue exclusion assay, and the percentage viability was calculated at different concentrations of the drug. The results demonstrated a shift in the IC50 towards a higher concentration (Table 3) in the presence of collagen when compared to control conditions. This suggests that the presence of collagen makes the cells more tolerant to chemotherapeutic drugs (Figure 2).

**Table 3:** IC50 of docetaxel across prostate cancer cell lines. The table shows the IC50 of docetaxel across different prostate cancer cell lines cultured either on a plain or a collagen-coated surface.

| **Cell line** | **Plain surface** | **Collagen-coated surface** |
| --- | --- | --- |
| LNCaP | 1.nM | 2.2nM |
| DU145 | 2.4nM | 3.1nM |
| PC3 | 2.4nM | 3.4nM |

**Figure 2:**
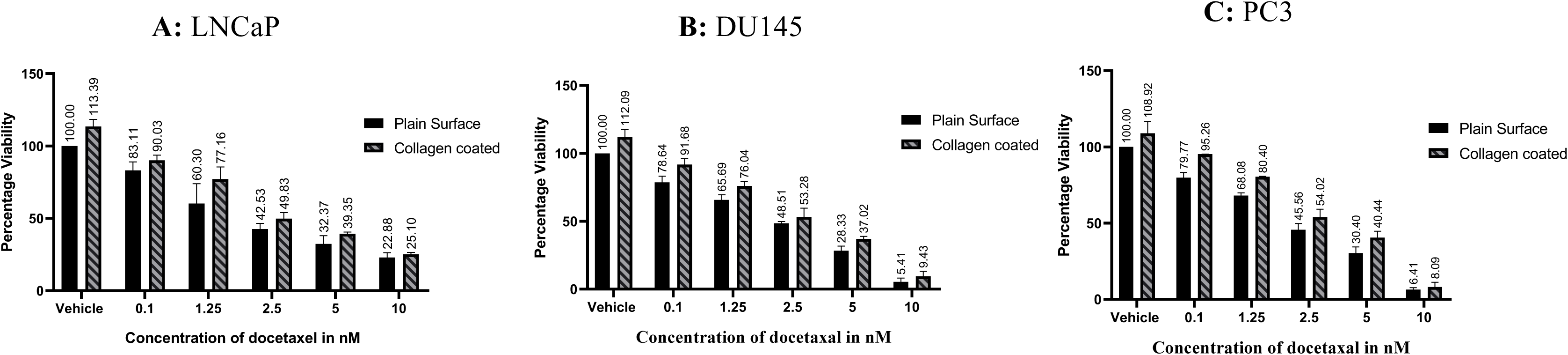
Determination of inhibitory concentration of docetaxel in prostate cancer cell lines cultured either on a plain surface or collagen-coated. Prostate cancer cell lines were cultured either on plain or collagen-coated surface of 24-well plates and treated with varying concentrations of docetaxel for 72hrs. Cell viability was assessed using the trypan blue exclusion assay. Each concentration was tested in duplicates, and the experiments were independently repeated three times.

### Effect of collagen on the migratory ability of cells

LNCaP, DU145, and PC3 cells were grown to confluence on either pre-coated collagen or plain 60mm dishes. A wound was created in the cell monolayer, and the width of the wound was measured every 4 hours. The rate of migration was calculated. No change in rate of migration was observed in the LNCaP cells cultured on the collagen-coated surfaces when compared to cells cultured on plain surfaces. An increased rate of migration was observed in cells grown on collagen-coated dishes compared to those on uncoated dishes in DU145 and PC3 cells (Table 4, Figure 3).

**Table 4:** Rate of migration of prostate cancer cell line cultured on collagen-coated or plain surfaces. The list shows the name of the cell lines and the rate of migration of cells cultured on collagen-coated surfaces and plain surfaces

| Cell line | Cells cultured on the plain dish | Cells cultured on a pre-coated collagen dish |
| --- | --- | --- |
| LNCaP | 1.18μ/hr | 1.27μ/hr |
| PC3 | 2.85μ/hr | 3.4μ/hr |
| DU145 | 3.7μ/hr | 4.3μ/hr |

**Figure 3:**
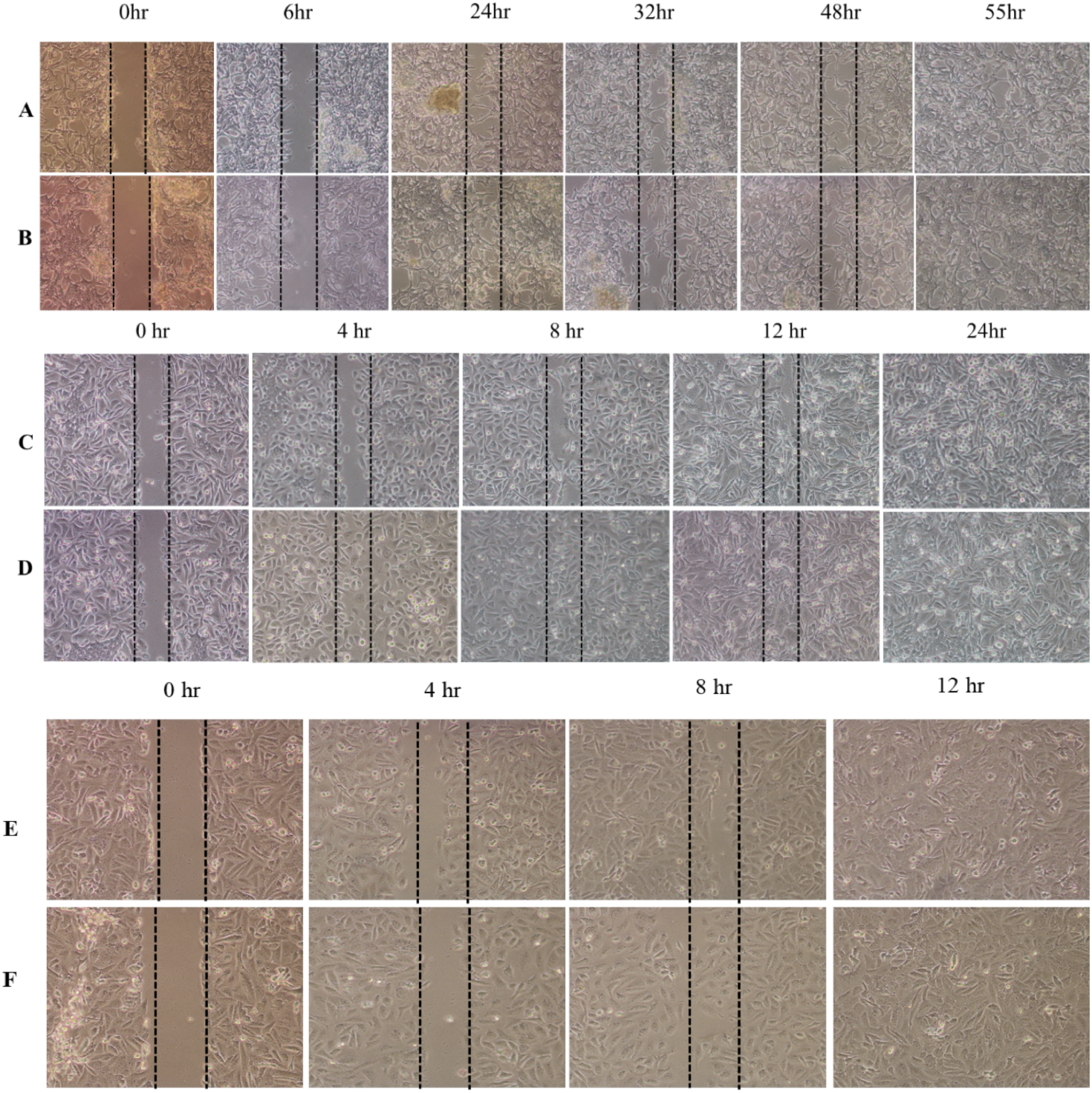
Representative images from an in vitro wound healing assay performed on prostate cancer cells cultured on either plain or collagen-coated surfaces. A total of 4X10^6^ cells/dish were plated either on plain or collagen-coated surfaces and allowed to grow until they reached the required confluency. A wound was created using a 200 µL micropipette tip. Images were captured post-wound to monitor cell migration. Scale bar: 100mm. A: LNCaP cells cultured on a plain surface. B: LNCaP cells cultured on a collagen-coated surface. C: DU145 cells cultured on a plain surface. D: DU145 cells cultured on a collagen-coated surface. E: PC3 cells cultured on a plain surface F: PC3 cells cultured on a collagen-coated surface

### Effect of collagen on stemness

LNCaP, DU145, and PC3 cells were grown to confluence on either pre-coated collagen or plain 60mm dishes. RNA was isolated and converted to cDNA. 20ng of RNA equivalent cDNA was used to perform stem cell marker (c-Myc, Klf4, Sox2, Oct4) qRT-PCR. The expression was normalized to the RPL35 gene. c-Myc showed reduced expression in LNCaP cells, whereas in DU145 and PC3 cells, there was an increased expression in response to collagen. Klf4 shows no significant change; Sox2 shows reduced expression in DU145 and PC3 cells; Oct4 shows increased expression in LNCaP and PC3 cells (Figure 4A, 4B, and 4C). Based on these results, the effect of collagen on stemness appears to be cell-specific. Results are summarized in Table 5.

**Table 5:** Stem cell markers expression status across different prostate cancer cell lines cultured either on a plain or collagen-coated surface. The list shows the status of stem cell markers across different prostate cancer cell lines

|  | **LNCaP** | **DU145** | **PC3** |
| --- | --- | --- | --- |
| Sox2 | No significant change | Reduced | Reduced |
| c-Myc | Reduced | Increased | Increased |
| Klf4 | No significant change | No significant change | No significant change |
| Oct-04 | Increased | No significant change | Increased |

**Figure 4:**
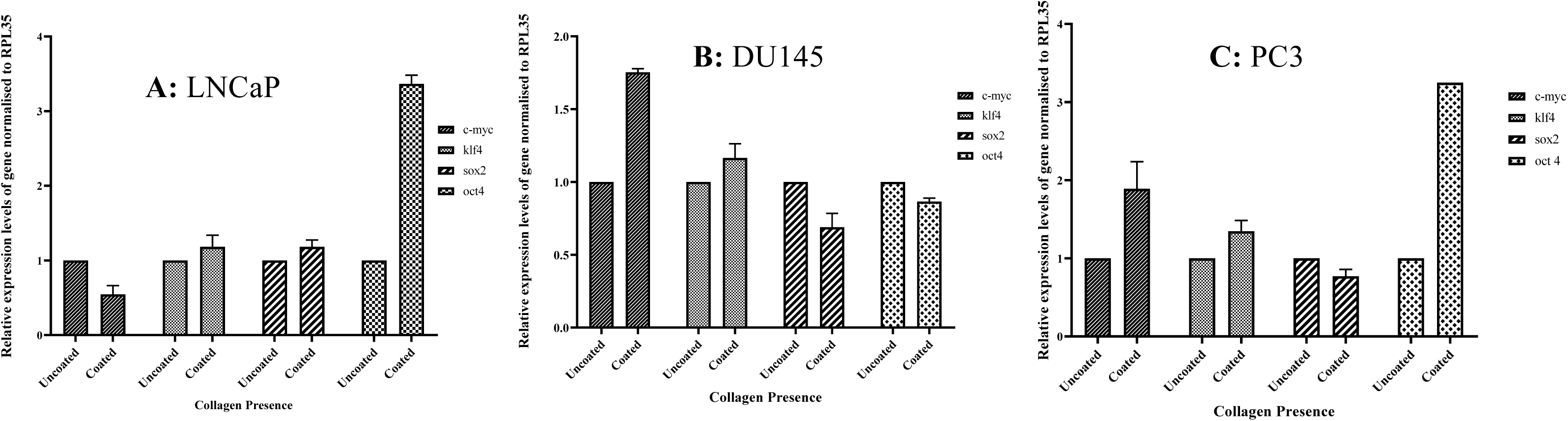
Expression of stem cell markers in prostate cancer cell lines cultured on a collagen-coated surface. The RNA was isolated from the prostate cancer cell lines cultured either on a plain or on a collagen-coated surface. The extracted RNA was converted into cDNA using reverse transcriptase and used as a template to perform qRT-PCR. The bar represents the relative expression levels of stem cell markers with respect to uncoated ± S.E.M., normalized to RPL35.

### Effect of collagen on androgen signaling

#### Effect of FAK on androgen signaling

Prostate-specific antigen (PSA) expression serves as a well-established direct measure of active AR signaling [21]. Clinically, PSA levels are routinely measured as a biochemical marker to assess prostate health. To assess the effect of collagen on AR signaling, qRT-PCR was performed to estimate the PSA gene expression in LNCaP cells, cultured on either collagen-coated or plain dishes. An increased expression of the PSA gene was observed in the RNA isolated from the collagen-coated dishes when compared with uncoated/plain dishes (Figure 5).

**Figure 5:**
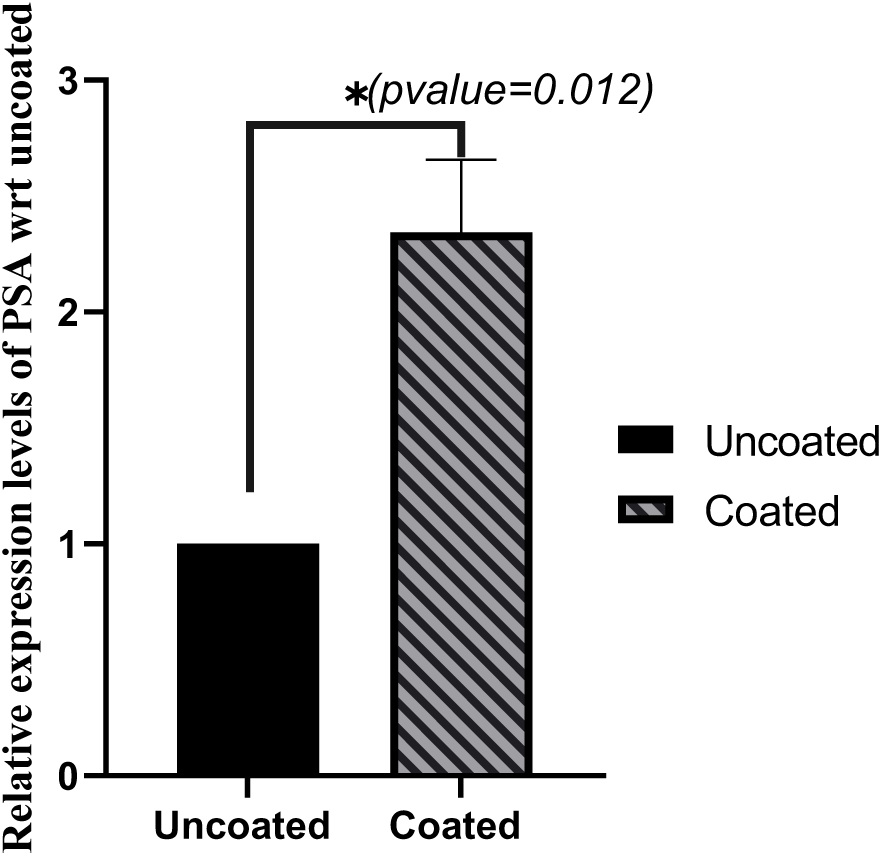
Expression of PSA in LNCaP cells cultured on a collagen-coated surface. RNA was isolated from the prostate cancer cell lines cultured either on a plain surface or on collagen-coated. The extracted RNA was converted into cDNA using reverse transcriptase and used as a template to perform qRT-PCR. The bar represents the relative expression levels of PSA with respect to uncoated ± S.E.M., normalized to RPL35.

Matrix remodeling is sensed by the integrins, leading to the activation of their cytoplasmic domain. This, in turn, triggers the activation and autophosphorylation of FAK. Activated FAK subsequently phosphorylates downstream proteins and initiates multiple signaling cascades, including androgen receptor signaling. To examine whether this elevated expression is due to FAK activation, a FAK inhibitor, PF 573228, was used. 5mM flutamide was used to create androgen-deprived conditions, and treated with or without 10nM PF 573228 for 48hrs. RNA was subsequently isolated, and PSA expression was estimated.

Flutamide decreased the expression of PSA in both uncoated and coated cells (Figure 6A). The FAK inhibitor significantly reduced PSA expression (Figure 6B). Furthermore, the combination of PF 573228 with flutamide also resulted in a significant decrease in PSA expression (Figure 6C). In all cases, the expression of PSA was higher in the collagen-coated condition compared to the control.

**Figure 6:**
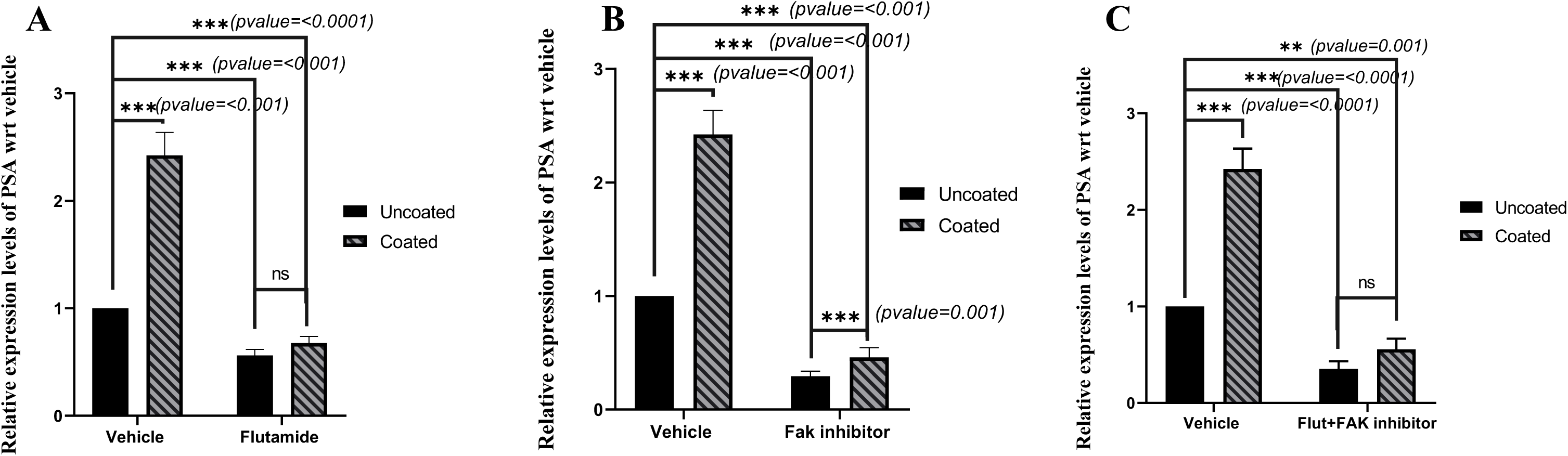
Expression of PSA in FAK inhibitor-treated LNCaP cells cultured on a collagen-coated surface. The RNA was isolated from the LNCaP cells cultured either on a plain or collagen-coated surface and treated +/- 10nM FAK inhibitor or +/- 5μM flutamide for 48hrs. The extracted RNA was converted into cDNA using reverse transcriptase and used as a template to perform qRT-PCR. The bar represents the relative expression levels of PSA with respect to vehicle ± S.E.M., normalized to RPL35.

### Effect of collagen on the EGFR pathway under androgen-deprived conditions

The role of the EGFR pathway in various cancers is well established. However, the contribution of this pathway to the development of androgen independence remains inadequately understood. To understand the role of TME-derived factors (in this case, collagen) on the EGFR pathway under androgen-deprived conditions, erlotinib, an EGFR pathway inhibitor, was utilized. Cells were plated either on the plain or collagen-coated surface, and RNA was collected after 48hrs of treatment. The expression of PSA was estimated using qRT-PCR. Both collagen-coated and uncoated conditions showed increased expression of PSA upon EGF treatment. Erlotinib treatment led to a reduction in PSA expression, which was partially restored by EGF co-treatment in both collagen-coated and uncoated (Figure 7A).

**Figure 7:**
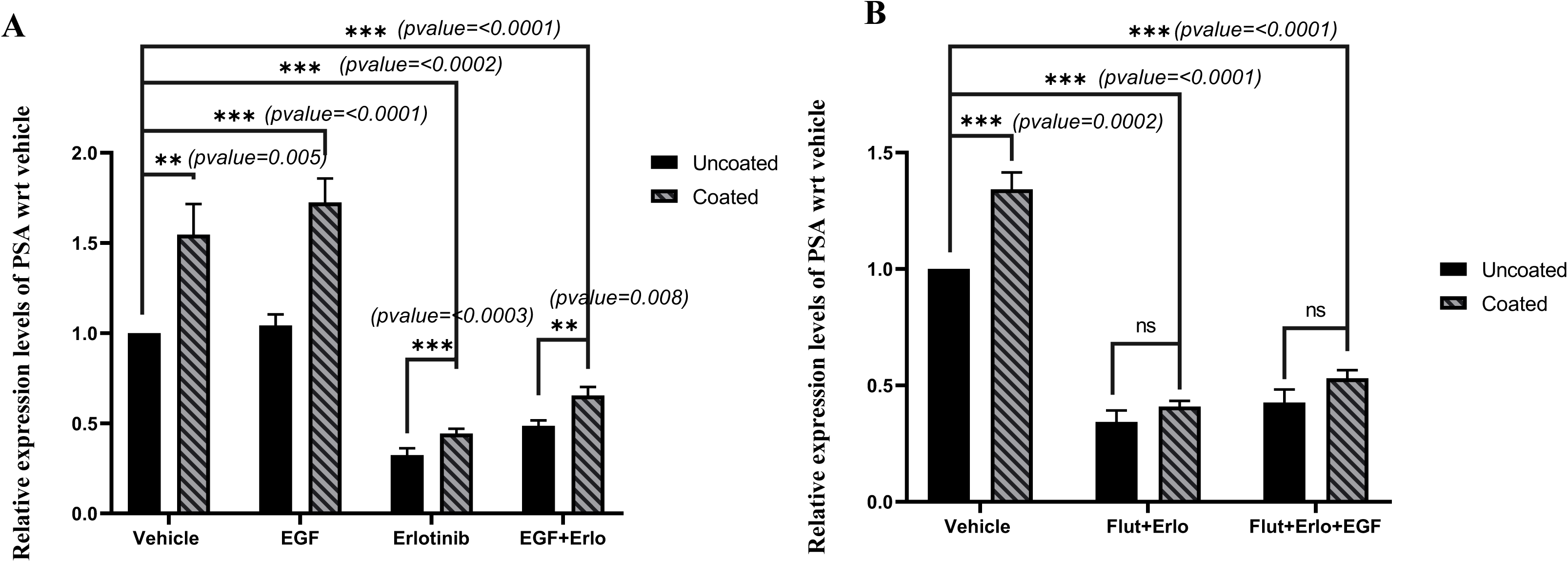
Expression of PSA in erlotinib-treated LNCaP cells cultured on a collagen-coated surface. RNA was isolated from LNCaP cells cultured either on a plain surface or collagen-coated and treated with +/-10nM EGF, +/- 5μM flutamide, or +/- 5μM erlotinib for 48hrs. The extracted RNA was converted into cDNA using reverse transcriptase and used as a template to perform qRT-PCR. The bar represents the relative expression levels of PSA with respect to vehicle ± S.E.M., normalized to RPL35.

Additionally, increased PSA expression was observed in collagen-coated and uncoated conditions upon EGF treatment, even in the presence of flutamide. However, simultaneous treatment with both flutamide and erlotinib resulted in decreased PSA expression in both the conditions (collagen-coated and uncoated) (Figure 7B). These findings suggest that simultaneously targeting the pathways contributing to androgen independence (EGFR in this case), in combination with AR-targeting agents, may offer greater potential to control tumor progression than AR-targeted therapy alone.

### Effect of collagen on cell proliferation

To evaluate the effect of collagen on cell proliferation, MTT was performed. The cells were plated either on collagen-coated or plain surfaces and treated with different combinations of inhibitors or activators. The cell viability was estimated using the MTT assay. Treatment with DHT increased the cell viability in both collagen-coated and uncoated (Figure 8A). Treatment with flutamide resulted in a reduction in cell viability. However, co-treatment with DHT partially restored cell viability in the presence of flutamide. A similar trend was observed with the FAK inhibitor; the combination of FAK inhibitor and DHT partially restored the cell viability as observed with flutamide and DHT (Figure 8B). Notably, while the combination of the FAK inhibitor and flutamide led to a further decrease in cell viability, the addition of DHT to this combination did not result in a significant rescue of cell viability (Figure 8C). Increased cell proliferation was observed on treatment with EGF in cells cultured on a collagen-coated surface. On treatment with erlotinib, cell viability was decreased and was partially restored on treatment with EGF (Figure 8D). A similar trend was observed in the cells treated with a combination of erlotinib, flutamide, and EGF (Figure 8E).

**Figure 8:**
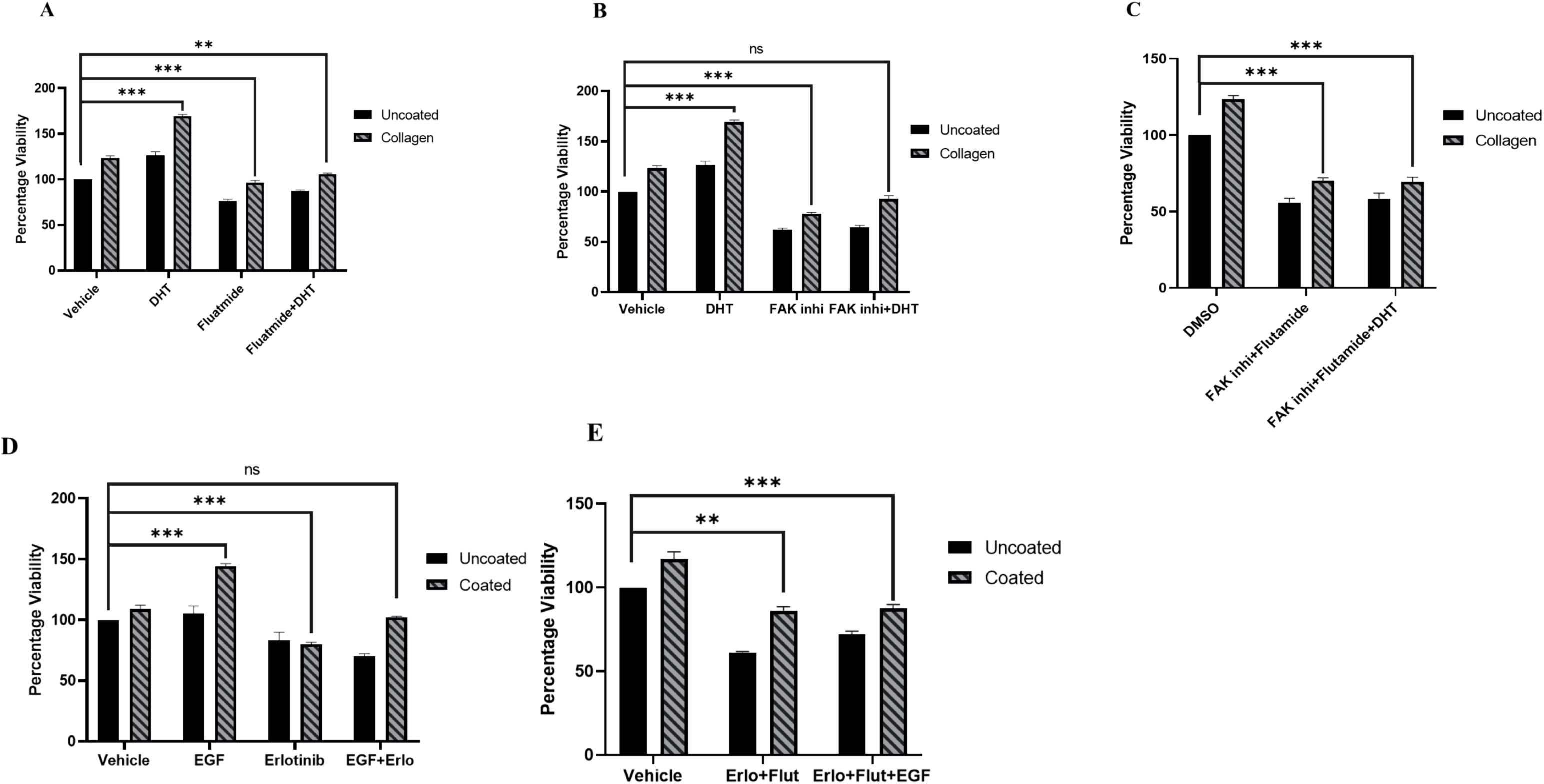
Effect of collagen on cell proliferation. A total of 5000 cells/well were treated with or without 10 nM DHT, +/- 5 μM erlotinib, +/- 5 μM flutamide, +/- 10 nM FAK inhibitor, +/- 10 nM EGF for 72 hrs. Cell proliferation was assessed using an MTT assay. The student t-test was performed to determine the significance between the groups. Each sample was in triplicate. The experiment was repeated twice.

However, in all these combination treatments, a slightly higher cell viability was observed in the cells grown on a collagen-coated surface compared to those on a plain surface, suggesting that collagen does contribute to enhanced cell proliferation.

## Discussion

The extracellular matrix is a highly dynamic structure that undergoes continuous deposition, remodeling, and degradation to maintain tissue homeostasis [5]. During cancer, ECM undergoes extensive alterations, directly influencing tumor cell proliferation, survival, invasion, migration, and metastasis. Increasing evidence indicates that ECM stiffening is a critical driver of tumor growth, invasion, and metastasis in prostate cancer. Malignant prostate tissues are almost 60% stiffer than benign prostate tissues [22]. ECM stiffening and its constituents are emerging as prognostic biomarkers for prostate cancer progression and metastasis. Increased collagen density and reorganization of collagen fibres correlate with the higher Gleason score, as more aligned collagen structures are observed in malignant biopsies compared to non-malignant tissue biopsies. These evidences indicate that ECM stiffening correlates with more aggressive forms of prostate cancer [22–24].

Several studies have investigated the impact of this ECM stiffness on cellular pathways. In castrate resistance prostate cancer (CRPC), ECM stiffness has been shown to activate key pathways such as EGFR, PI3K/AKT, and MAPK [25–27]. Notably, matrix stiffness induces the activation of multiple signaling pathways that overlap with those involved in both androgen-dependent and androgen-independent prostate cancer progression. These findings suggest that ECM stiffness contributes to the molecular mechanisms underlying prostate cancer aggressiveness and therapy resistance.

Collagen is one of the major components of the ECM. Various types of collagens are differentially expressed in cancerous vs non-cancerous tissues, for example, collagen XXII, and can be detected in urine. [17, 28]. A significant alteration in collagen metabolism has been observed in cancerous tissue compared to adjacent histologically benign prostatic tissue [18]. A study demonstrated that type 1 collagen serves as a primary substrate for PC3 cells, a prostate cancer cell line. Moreover, collagen type 1 was found to significantly enhance PC3 cell proliferation through activation of the PI3K pathway, suggesting that collagen may promote prostate cancer cell metastasis by increasing both cellular attachment and proliferation [29]. Additionally, type 1 collagen was shown to enhance the invasion capacity of the human prostate cancer cell line LNCaP, via α2β1 integrin and RhoC signaling. Treatment of LNCaP cells with a neutralizing antibody against α2β1 integrin decreased collagen-induced in vitro invasion, indicating that α2β1-mediated signaling facilitates collagen-driven invasion through the upregulation of RhoC. Collectively, these studies suggest that collagen plays a fundamental role in promoting prostate cancer metastasis [30].

However, limited studies have investigated the precise mechanisms by which collagen contributes to cancer progression, highlighting the importance of elucidating the role of collagen in cancer [31, 32]. Our study showcased that the collagen may activate/influence other signaling pathways. We hypothesize that matrix stiffness due to the deposition of TME-derived factors such as collagen may activate integrins and FAK, leading to the activation of non-genomic signaling /genomic signaling of AR, thereby contributing to androgen independence. Additionally, matrix stiffness may also influence the EGFR signaling cascades. The activated EGFR pathway cascade (MAPK, PI3K/AKT) may, in turn, induce the PSA gene transcription through its interaction with the PSA upstream enhancer containing AP-1 binding sites (-4420) [33] (Figure 9). Also, the deposition of collagen or other TME derivatives may influence the effective intake of drugs by tumor cells. Therefore, targeting some of these factors and associated signalling, such as FAK signalling, may be a better therapeutic strategy to manage prostate cancer. particularly to prolong the duration of the androgen-sensitive phase in patients.

**Figure 9:**
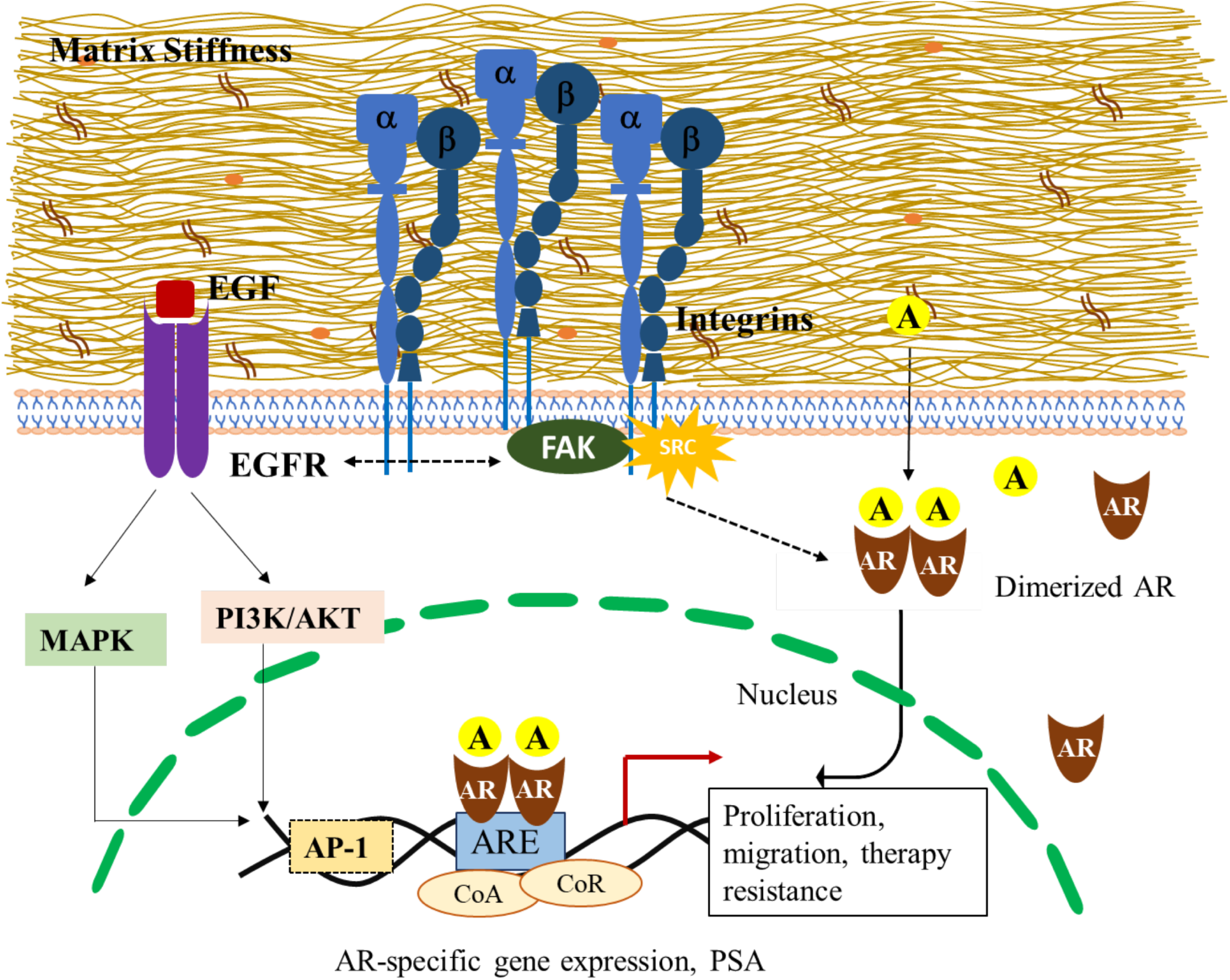
Schematic representation of signalling pathways activated by ECM stiffness.

